# Bioinformatic investigation of discordant sequence data for SARS-CoV-2: insights for robust genomic analysis during pandemic surveillance

**DOI:** 10.1101/2023.02.01.526694

**Authors:** Sara E. Zufan, Katherine A. Lau, Angela Donald, Tuyet Hoang, Charles S.P. Foster, Chisha Sikazwe, Torsten Theis, William D. Rawlinson, Susan A. Ballard, Timothy P. Stinear, Benjamin P. Howden, Amy V. Jennison, Torsten Seemann

## Abstract

The COVID-19 pandemic has necessitated the rapid development and implementation of whole genome sequencing (WGS) and bioinformatic methods for managing the pandemic. However, variability in methods and capabilities between laboratories has posed challenges in ensuring data accuracy. A national working group comprising 18 laboratory scientists and bioinformaticians from Australia and New Zealand was formed to improve data concordance across public health laboratories (PHLs). One effort, presented in this study, sought to understand the impact of methodology on consensus genome concordance and interpretation. Data were retrospectively obtained from the 2021 Royal College of Pathologists of Australasia Quality Assurance Programs (RCPAQAP) SARS-CoV-2 WGS proficiency testing program (PTP), which included 11 participating Australian laboratories. The submitted consensus genomes and reads from eight contrived specimen were investigated, focusing on discordant sequence data, and findings were presented to the working group to inform best practices. Despite using a variety of laboratory and bioinformatic methods for SARS-CoV-2 WGS, participants largely produced concordant genomes. Two participants returned five discordant sites in a high Ct replicate which could be resolved with reasonable bioinformatic quality thresholds. We noted ten discrepancies in genome assessment that arose from nucleotide heterogeneity at three different sites in three cell-culture derived control specimen. While these sites were ultimately accurate after considering the participants’ bioinformatic parameters, it presented an interesting challenge for developing standards to account for intrahost single nucleotide variation (iSNV). Observed differences had little to no impact on key surveillance metrics, lineage assignment and phylogenetic clustering, while genome coverage <90% affected both. We recommend PHLs bioinformatically generate two consensus genomes with and without ambiguity thresholds for quality control and downstream analysis, respectively, and adhere to a minimum 90% genome coverage threshold for inclusion in surveillance interpretations. We also suggest additional PTP assessment criteria, including primer efficiency, detection of iSNVs, and minimum genome coverage of 90%. This study underscores the importance of multidisciplinary national working groups in informing guidelines in real time for bioinformatic quality acceptance criteria. It demonstrates the potential for enhancing public health responses through improved data concordance and quality control in SARS-CoV-2 genomic analysis during pandemic surveillance.

**Data summary:** The authors confirm all supporting data, code and protocols have been provided within the article or through supplementary data files.

**Impact statement:** Amidst the COVID-19 pandemic, a unique collaboration between a national multidisciplinary working group and a quality assurance program facilitated ongoing development of standardized quality control criteria and analysis methods for high-quality SARS-CoV-2 genomic approaches across Australia. With this article, we shed light on the robustness of amplicon sequencing and analysis methods to produce highly concordant genomes, while also presenting additional assessment criteria to guide laboratories in identifying areas for improvement. Insights from this nationwide collaboration underscore the need for real-time knowledge-sharing and iterative refinements to quality standards, particularly as situations and methods evolve during a pandemic. While the spotlight is on SARS-CoV-2, the analyses and findings have universal implications for genomic surveillance during infectious disease outbreaks. As WGS becomes increasingly central in outbreak surveillance, continuous evaluation and collaboration, like that described here, are vital to ensure data accuracy and inform future public health responses.

## Introduction

The global scientific community responded to the coronavirus disease (COVID-19) pandemic with the rapid development of laboratory and bioinformatic data analysis methods for whole genome sequencing (WGS) of severe acute respiratory syndrome coronavirus 2 (SARS-CoV-2). These methods have been widely adopted, with more than 45 countries conducting routine genomic surveillance (1). Genomic data have been used to inform the local and national pandemic response. Applications range from monitoring the distribution and emergence of variants of concern (VOC) to supplementing contact tracing efforts (2). These efforts have been supported by an open-access framework for the wide-scale sharing of data and resources, including Global Initiative on Sharing All Influenza Data (GISAID), Nextstrain, and protocols.io, among others. By May 2020, problematic sequence data attributed to contamination or sequencing and bioinformatic errors had been detected in the public domain, highlighting the need for quality control and remediation strategies to improve consensus genome accuracy (3).

Amplicon sequencing has been the main method of SARS-CoV-2 WGS, with bioinformatic methods dependent on the platform (4,5). Previous interlaboratory and interprotocol comparative studies of SARS-CoV-2 WGS have found differences in single nucleotide variants (SNV) during the assessment of wet and dry methodologies (6,7). The appropriate selection of bioinformatic workflows and parameter thresholds is key to the accuracy of genomic data and varies depending on the sequencing approach and technology. While selection varies, some commonly accepted thresholds include: read depth (>=10 for Illumina, >=20 for Nanopore), phred score (>25), and genome coverage >=90% (8). Discrepancies at SARS-CoV-2 mutation sites can affect the interpretation of genomic metrics of clinical and public health importance, such as PANGO lineage classification, phylogenetic placement, or genomic epidemiological clustering (9–11). As such, bioinformatic quality control processes ensure that methodological variability yields consistent results.

Australia and New Zealand have been at the forefront in the use of fine-scale genomic data to manage their COVID-19 responses. Genomic data are shared via AusTrakka to provide context for local and interstate transmission. AusTrakka is a national real-time pathogen genomics surveillance platform for public health laboratories that facilitates analytical harmonisation and national reporting as well as provides equitable access to computational resources and expertise (12). Australia’s high case sequencing rate, combined with multijurisdictional data sharing, resulted in genomic epidemiological interpretations with as little as one single nucleotide variant (SNV). As is common in many countries, each jurisdiction has a unique laboratory and bioinformatic workflow to suit its facility.

In July 2020, the Medical Research Future Fund (MRFF) COVID-19 Quality Control Working Group (WG), supported by the Communicable Diseases Genomics Network (CDGN), initiated a concerted effort to harness genomic sequencing for an enhanced response to the COVID-19 pandemic. Comprised of 18 public health laboratorians and bioinformaticians from Australia, the WG’s primary objective was to establish a framework that ensures laboratory and bioinformatic standards and quality control, permitting procedural adaptability yet ensuring consistent SARS-CoV-2 genomic data. Continuous communication among laboratory technicians and bioinformaticians was maintained via monthly meetings and regular online discussions via Slack, facilitating protocol optimisation and addressing potential sequencing and analysis challenges in real time.

The WG aimed to enhance interlaboratory data concordance but faced challenges in coordinating a national validation study during pandemic surges. Consequently, the WG procured SARS-CoV-2 WGS proficiency testing program (PTP) data from the Royal College of Pathologists of Australasia Quality Assurance Programs (RCPAQAP) for retrospective, in-depth bioinformatic analyses. In the study presented here, PTP data was used to characterise discordant SNVs and investigate bioinformatic quality control standards for remediation. Submitted consensus genomes were used to evaluate the effect of discordant SNVs on key surveillance metrics, lineage assignment and phylogenetic clustering, while submitted read data were used to investigate discordant sites. Finally, we reflect on the importance of multidisciplinary national working groups in informing guidelines in real time for bioinformatic quality acceptance criteria under competing demands of a pandemic.

## Methods

### Data Source: 2021 RCPAQAP SARS-CoV-2 WGS PTP

Data were obtained from the 2021 RCPAQAP SARS-CoV-2 WGS PTP, sample characteristics and methods have previously been described (13). Briefly, viral RNA extracted from five unique isolates of SARS-CoV-2 isolates were each diluted in 0.5 ml of nuclease-free water to assess the participant’s ability to sequence different variants and viral titers.

Contrived samples (N=8) were sent to 11 participating laboratories for WGS using the laboratory’s standard procedure. The raw reads of sequence data without pre-processing or trimming and the derived consensus sequence data, as submitted by the participants, were assessed based on three quality metrics (>50% genome coverage, lineage assignment, and >95% accuracy), as decided with input from CDGN Genomics Implementation and Bioinformatics Working Groups and previously described by Lau et al (2022) (13). The participants also completed a questionnaire documenting laboratory and bioinformatic methods performed and self-reported genomic characteristics.

Deidentified data were provided to the WG for the principal aim of defining QC criteria and data analysis methods for standardising high-quality SARS-CoV-2 WGS, independent of the PTP. One participant belonging to the WG inadvertently omitted consensus genomes from their submission to the PTP and provided them voluntarily for this study. Herein, we refer to the participants by their laboratory number (LB01 – LB11) and the samples by their biological specimen number (BS01 – BS08) (Table 1).

**Table 1.** Characteristics of contrived specimen prepared for the PTP.

| Sample ID | Isolate (GISAID ID) | Average Ct (E-gene NAT) | Lineage |
| --- | --- | --- | --- |
| BS01 | EPI_ISL_406844 | 16.77 | B |
| BS02 | EPI_ISL_419750 | 16.95 | B.1 |
| BS07 |  | 24.14 |  |
| BS08 |  | 27.48 |  |
| BS03 | EPI_ISL_480701 | 17.28 | B.1.1.136 |
| BS04 | EPI_ISL_519314 | 17.31 | D.2 |
| BS05 | EPI_ISL_563416 | 17.29 | D.2 |
| BS06 | Negative Control | Not detected | NA |

### Evaluation of consensus genomes

For the assessment of consensus genome quality, genome coverage and SNVs were characterised. To discern the influence of these quality parameters on critical surveillance metrics, consensus genomes were subjected to PANGO lineage assignment and phylogenetic clustering.

#### Characterisation of consensus genome quality

Genome coverage was calculated as the percentage of non-N alleles in a consensus genome relative to the SARS-CoV-2 reference genome (Wuhan-Hu-1; GenBank MN908947.3).

Consensus genome positions where observed alleles matched expected alleles were characterised as ‘concordant’, ‘discordant’ if the consensus allele did not match the expected allele, ‘missing’ if there was an N or gap at an expected SNV position of the SNV, or ‘ambiguous’ if any position contained an IUPAC ambiguity code.

#### Lineage assignment

The consensus genomes submitted in FASTA format were used to obtain genome coverage and PANGO lineages. Lineages were assigned to the samples using Pangolin v.2.3.3 with pangoLEARN 2021-21-02, versions available at the time of the PTP. Discordant, missed, and ambiguous SNVs relative to expected SNVs defined by the PTP were identified using Nextclade v.2.8.0 relative to expected SNVs defined by the PTP (Figure 1).

**Figure 1.**
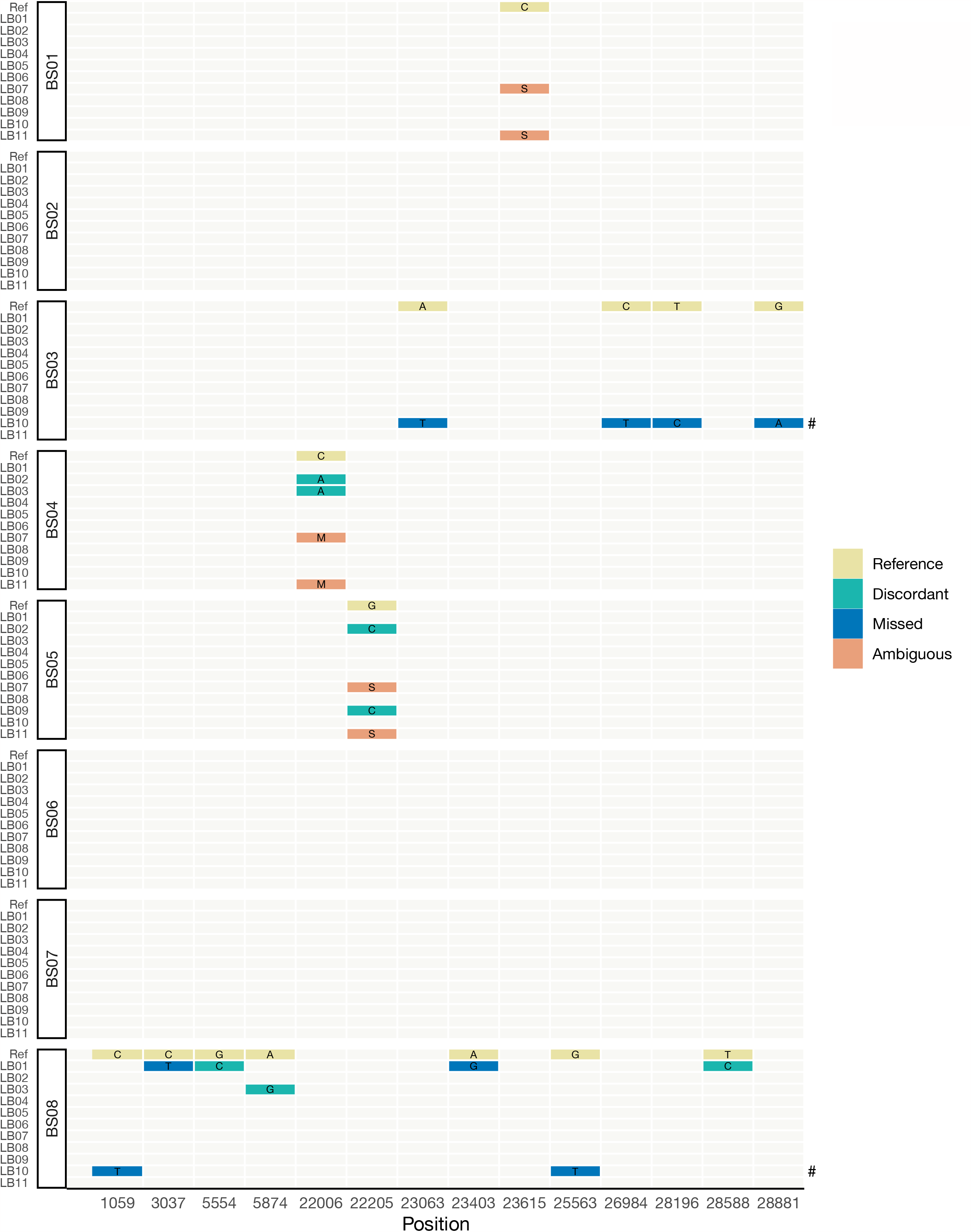
Discordant SNVs observed in participant consensus genomes submitted. Reference indicates the expected allele of the sample isolate reference sequence. Discordant SNVs are those where the observed participant allele differs from the expected reference allele. Missed SNVs are a subset of discordant SNVs in which a participant called an allele different from the expected SNV. Ambiguous SNVs are another subset of discordant SNVs in which the participant called an IUPAC ambiguity code where a standard base (ATGC) was expected (S = C or G; M = A or C). According to the criteria used in the PTP, the absence of C241T was considered concordant, and thus omitted from this figure if present. No annotation implies concordance. #LB10 had additional discordant SNVs for samples BS03 (N=51) and BS08 (N=33) that are not listed here.

#### Phylogenetic clustering

A phylogenetic tree was constructed using RAxML-NG v.1.1 with a GTR+G4 model, starting trees from FastTree v.2.1.11 with 1,000 bootstraps, and alignment column weight using gotree goalign compress v.0.4.2 (14). BS03 and BS08 from LB10 were omitted from the final input alignment due to disproportionately long branches that negatively affected tree readability.

### Bioinformatic investigation of sequencing reads

To assess participant methods from the PTP survey for bioinformatic investigation, it was found that responses could not be consistently reconstructed. Consequently, a standardised workflow was adopted for Illumina and Nanopore data to investigate base pair attributes at discordant sites and amongst amplicons. Both trimmed and untrimmed reads were investigated

#### Generating read pileups

FASTQ reads were aligned to the SARS-CoV-2 reference employing minimap2 v. 2.24-r1122, using the options -ax map-ont for Nanopore or -ax sr for Illumina reads. Relevant amplicon primers (Supplementary Table 1) were trimmed from the aligned reads via Samtools ampliconclip v.1.16.1, employing options --both ends --strand --soft-clip -u.

#### Obtaining base pair attributes

Base pair attributes of the total aligned reads were quantified using bam-readcount v.1.0.1 with default settings and subsequently interpreted using the parse_brc.py script (15). Trimmed reads were processed similarly with option -b 20 to discard low-quality reads. The read depth for each amplicon was determined using mosdepth v.0.3.3, utilizing the corresponding amplicon insert regions as the input for the bam file.

### Investigating iSNVs in primary isolates

Investigation of base pair attributes revealed heterozygous sites in three specimen recovered by all participants. To confirm the presence of heterozygous alleles in the primary isolates used to create the biological specimen, likely to be iSNVs, we retrieved their respective sequencing reads. Isolates were amplicon sequenced using ARTIC V3 primers and sequenced on an Illumina iSeq. Base pair attributes were obtained as described earlier.

## Results

### Summary of reported methodology

Participant workflows varied for laboratory and bioinformatic methodologies (Supplementary Tables 1-2). Most participants (N=9/11) performed multiplex amplicon sequencing, with seven using the ARTIC V3 primer scheme (16,17), another using a modified ARTIC V3 primer scheme to create 1 kbp amplicons, and one using the ‘Midnight’ scheme (4,18). Two participants used a long-pooled amplicon approach, ‘JSE’, with primers that produce 2.5 kbp amplicons (19,20). Most of the participants used an Illumina sequencing platform (N=8/11), with 6/8 using ARTIC V3 primers. All participants using Nanopore technology used the artic-ncov2019 pipeline (https://github.com/artic-network/artic-ncov2019). Amongst Illumina users, four used an in-house analysis pipeline, two used a mix of command line and commercial software, and two used solely commercial software (CLC Genomics Workbench).

### Characteristics of consensus genome quality

#### Genome coverage and contamination

Mean genome coverage for all consensus sequences submitted of SARS-CoV-2 positive samples was 95.98%. Mean coverage of BS08 was lower overall (93.75%) due to a high cycle threshold value (Ct=27.48) (13). LB05 and LB09 had consistent coverage <99%, suggesting dropout of the amplicon(s). While most participants detected few to no SARS-CoV-2 reads in the negative control BS06, LB03 and LB05 recovered 11.0% and 4.5% of the genome, respectively, and LB10 recovered 79.2% (Supplementary Figure 1).

#### Discordant SNVs

Based on the consensus genome alone, 4/11 participants (LB04-6,8) submitted 100% concordant genomes, which included all Nanopore users (LB04-6) that used different primer schemes (JSE, V3, Midnight) (Figure 1). Two participants (LB07,11) were excluded from 100% concordance solely due to ambiguous bases. The remaining 5/11 participants had one or more discordant SNV. Except for LB10, discordant SNVs per participant ranged from one to four. In the consensus genomes submitted by LB10 for BS03 and BS08, there were 55 and 35 SNVs, respectively. Excluding ambiguity calls and LB10, participants generated 100% concordant genomes for BS01-3 and BS07.

### Impact on key surveillance metrics

#### Lineage assignment

PANGO lineage assignments were largely tolerant of missing data and false SNVs, with 95% of consensus genomes submitted correctly designated at the highest taxonomic level (Figure 2a). Because PANGOLIN classification is hierarchical, genomes with missing data can be expected to have parent lineage assignments, such as BS03 from LB10 (21). BS08 samples from LB01 and LB10 were probably designated as a higher, incorrect lineage due to a combination of low genome coverage and false SNVs.

**Figure 2.**
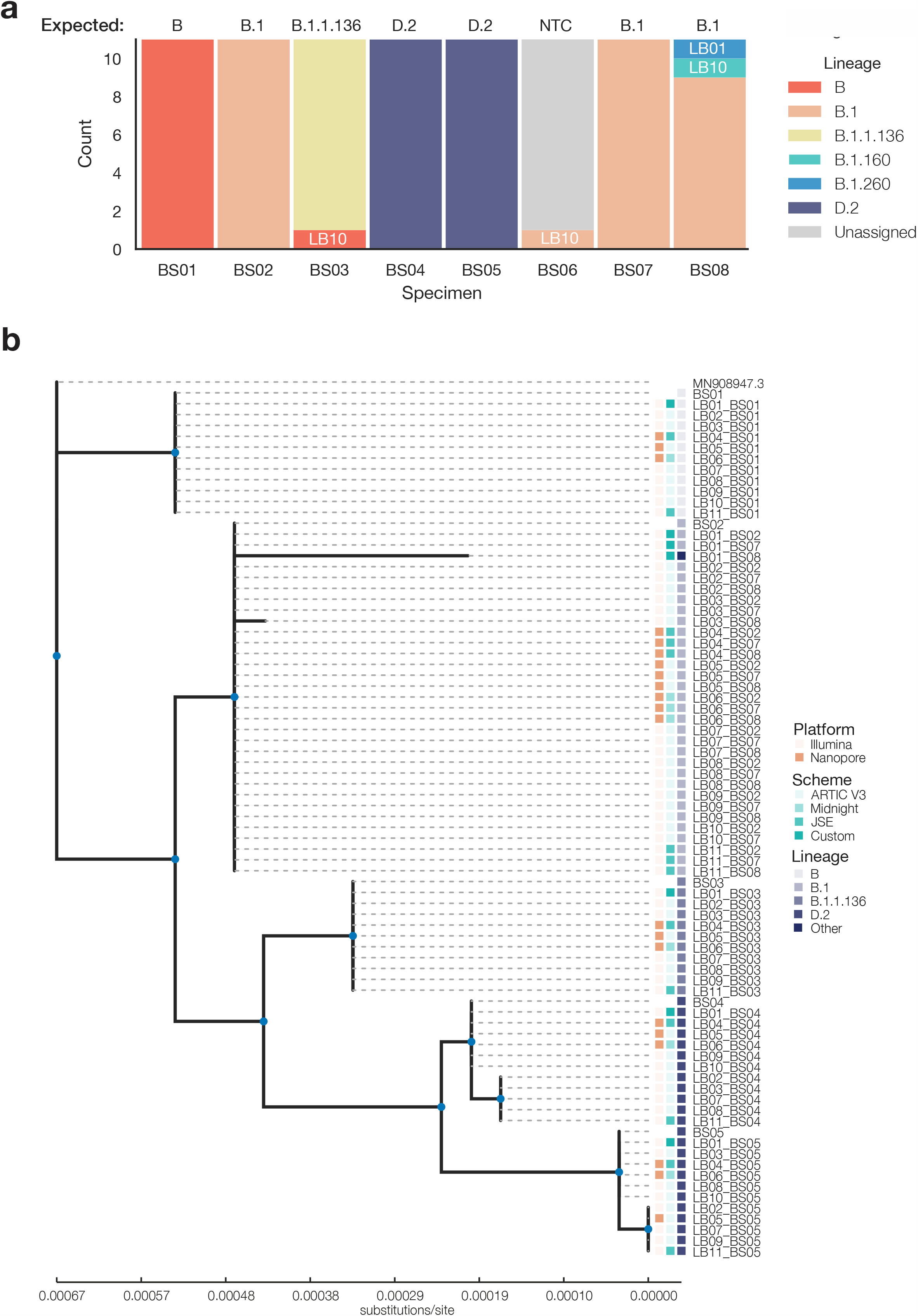
Key surveillance metrics derived from consensus genomes submitted. A) Pangolin lineage assignments from consensus genomes submitted. Participant ID annotated where discordant. b) Maximum likelihood tree of consensus genomes submitted, excludes genomes <50% coverage; and BS03 and BS08 for LB10. Nodes with teal markers indicate branch support > 80%. The heatmap denotes the sequencing platform, primer scheme, and PANGO lineage called for each tip. Phylogenetic tree visualised with toytree v1.0 (https://github.com/eaton-lab/toytree).

#### Phylogenetic clustering

In general, the consensus genomes submitted were placed in their expected phylogenetic clades (Figure 2b). BS03 and BS08 from LB10 were omitted from the input alignment as the relatively high number of false mutations resulted in long branches, making it difficult to interpret the tree. Among the remaining 75 genomes, 84% were placed within their expected clade. Subclades formed with low bootstrap support in BS04 and BS05 which included specimens with ambiguity codes, those that had a majority allele different from the isolate reference, and one specimen that was concordant with the basal clade. BS08 samples from LB01 and LB03 formed longer branches within the expected clade, with the former having the longest branch due to lower genome coverage (63.0% vs. 96.7%) and more discordant SNVs (N=4 vs. N=2).

### Bioinformatic investigation of sequencing reads

#### Amplicon dropouts

Per-amplicon read depth revealed LB05 had two or more amplicon dropouts across all samples, generally in amplicons 64 and 70. Similarly, LB09 failed to amplify amplicons 1 and 98 in all samples.

#### Base pair attributes at heterozygous sites

Inspection of read pileups at discordant positions associated with ambiguity codes found that all participants had sequenced isolates with allele heterogeneity in BS01, BS04, and BS05 (Figure 3a). Therefore, the allele called at the position depended upon a) the allele mixture sequenced; b) the variant calling threshold; and c) whether ambiguity codes were used in variant calling. Taking this into account, six participants had 100% concordant genomes, with an additional two having appropriate ambiguity codes.

**Figure 3.**
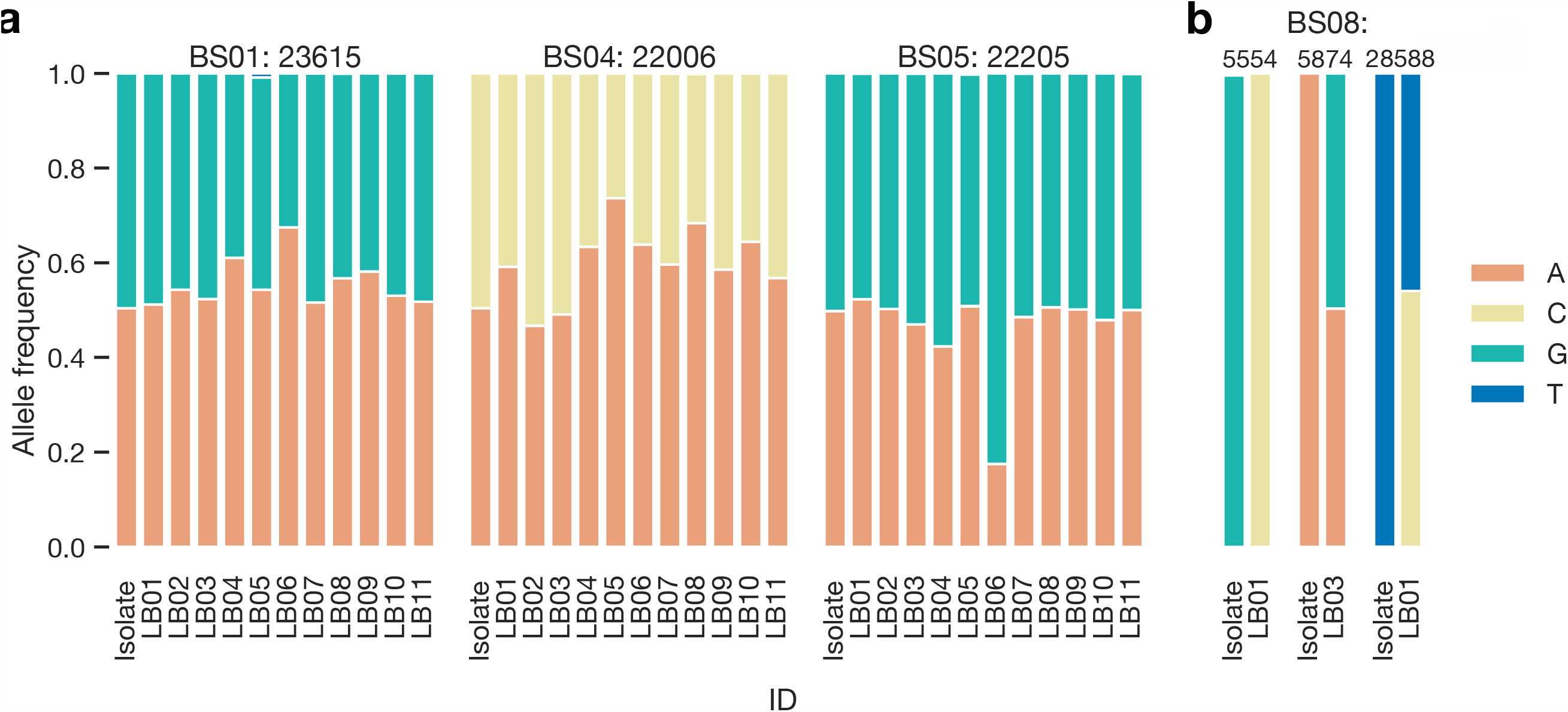
Allele distribution at discordant positions. a) Allele frequencies at positions containing ambiguity in consensus genomes submitted to the PTP and the primary isolate used to create contrived specimen. b) Allele frequencies at remaining discordant positions in both participant and primary isolate reads.

#### Base pair attributes at discordant sites

LB01 and LB03 yielded discordant or missed SNVs in BS08, the high-Ct sample. At nucleotide (nt) positions 25,588 and 5,574 in LB01 and LB03, respectively, we observe an allele mixture where the discordant allele is the major allele according to the variant calling thresholds of the participant. We investigated the allele frequency at identical sites in the remaining samples and found them homogeneous. Finally, at nt position 5,455 in the sample of LB01, the discordant allele was present at a frequency of 1 (Figure 3b). Given that this false SNV was present in 100% of the reads and was not detected in the other matched specimen, it may be the product of chimeric PCR amplification (7).

#### Analysis of problematic SNVs in LB10

LB10 had the highest number of discordant SNVs, with 55 and 35 in BS03 and BS08, respectively. Given the high recovery of the SARS-CoV-2 genome in the negative control of LB10 and the high number of total reads, laboratory contamination was suspected. However, an inspection of the read pileup found a disproportionately high read depth among several regions and found that the concordant allele was the major allele at discordant positions in many instances.

In BS03, there was overamplification of amplicons 2, 3 and 4 with a mean read depth of 43866, 302975 and 47687, respectively. The mean depth across all other amplicons was 33.6. In BS08, the mean amplicon depth was 18.2. However, the read depth in the 3’UTR was 5910.7. Inspection of the read pileup in this region revealed an abundance of poly-A reads.All LB10 samples had similar read mapping characteristics in the 3’ UTR, possibly suggesting poly-A carryover from cDNA synthesis.

When investigating the distribution of alleles at discordant positions, in many cases, the discordant allele was present at a depth of 1 or 0 (LB10-BS03: 78.2%; LB10-BS08: 91.4%) (Supplementary Figure 2). Where the expected concordant allele was present in all or a majority of reads at a discordant site the mean frequency was 0.95 (median: 1.0; IQR: 0.05) and 0.96 (median: 1.0; IQR: 0.0) in LB10-BS03 and LB10-BS08, respectively. In fewer cases (LB10-BS03: 14.5%; LB10-BS08: 5.71%), the discordant allele was present in greater frequency than the concordant allele where the discordant allele had a mean frequency of 0.79 (median: 0.81; IQR: 0.23) and 0.62 (median: 0.62; IQR: 0.12) in LB10-BS03 and LB10-BS08, respectively. These observations of higher discordant allele frequency, along with the high presence of SARS-CoV-2 reads in the negative control specimen, may suggest these sites are the result of contamination.

We could not determine this participant’s specific bioinformatic methods or parameters from their questionnaire responses to further investigate how variants were being called (Supplementary Table 2). However, we can infer that the minimum depth threshold for variant calling was below 10, a commonly accepted minimum criterion for Illumina amplicon sequence data (8).

## Discussion

Here, we report findings from an in-depth retrospective analysis of discordant SNVs using consensus genomes, raw reads, and the method questionnaire from a national SARS-CoV-2 WGS PTP. Despite diverse laboratory and bioinformatic methods, the data yielded largely consistent genomes, with discrepancies often resolved using robust bioinformatic thresholds. These discrepancies minimally affected key surveillance metrics. Recognizing that consensus genomes are generally accurate, certain challenges like low read depths and non-uniform amplicon sequencing coverage arise. As laboratory and bioinformatic techniques have advanced throughout the pandemic, there’s a pressing need to continually refine SARS-CoV-2 WGS quality standards, which was facilitated here by the WG’s collaborative efforts.

### Flexibility of amplicon sequencing analyses

We find that consensus genomes generated with bioinformatic parameters at or above the commonly accepted minimum thresholds were highly accurate. It should be noted that two participants (LB03 and LB10) generated consensus genomes using read depth <10, the commonly accepted minimum threshold, with mixed results. LB03 used a depth of 3, resulting in 100% concordant genomes when including ambiguity. It may be that LB10 used a depth of 1, resulting in 6/8 consensus genomes with 100% concordance and two (BS03, BS08) with >40 discordant SNVs. One concern regarding variant calling with low read depth is false SNVs due to contamination, as observed here for a small number of false SNVs in LB10-BS03 and LB10-BS08. This can be ameliorated by monitoring reads in a negative sequencing control or a consensus sequence control to determine if a read set should use a higher depth threshold or fail quality control. The latter approach is more bioinformatically challenging to extract and interpret allele frequencies.

All Nanopore users (LB04-6) generated 100% concordant genomes despite each using different primer schemes and lower read quality compared to Illumina users, as typically observed. Unlike Illumina users who used custom analysis scripts or commercial software, all Nanopore users employed the artic-ncov2019 pipeline (https://github.com/artic-network/artic-ncov2019. The robustness of the pipeline is unsurprising as it was designed especially for amplicon sequencing with Nanopore. While we could not reconstruct the analyses from the methodology questionnaire, our findings nonetheless suggest further coordination between Illumina users is needed to troubleshoot problematic thresholds or standardise an analysis workflow.

### Considerations for assessing amplicon sequencing data

For PHLs already engaged in WGS quality control, they have typically had more experience with methods designed for prokaryotic pathogens in which shotgun sequencing of isolates is performed (22–24). The characteristics of amplicon sequencing differ in several ways in which technical bioinformatic analysis can support laboratory optimisation. First, PCR efficiency often results in non-uniform coverage across the SARS-CoV-2 genome. Although genome coverage is an important quality indicator for unbiased sequencing approaches, it can be misleading when a small number of amplicons are overamplified or poly-A is carried through the library preparation and sequenced, as observed here with LB10. Instead, monitoring the mean read depth per amplicon would help participants troubleshoot amplicon drop outs, a common issue with this method that requires optimisation (25–27). Second, the proximity of discordant SNVs to primers can indicate bioinformatic trimming errors. It has previously been observed that CLC Genomics

Workbench (Qiagen, https://www.qiagenbioinformatics.com/products/clc-genomics-workbench, accessed 03 Jan 2023) has a lower sensitivity to trimming partial primers and calling SNVs in primer regions, compared to iVar (7). Although we were unable to replicate the results of LB10 using CLC 22 (*Identify ARTIC V3 SARS-CoV-2 Low-frequency and Shared Variants (Illumina) Workflow*) we did observe that false SNVs occurred near primer sites, which could signal the need for bioinformatic troubleshooting by the participant.

### Recommendations for assessing heterozygous sites

Similar to Foster et al. (2022), we primarily observed interlaboratory differences between matched samples when iSNVs, or heterozygous sites, were present (6). As in the assessments performed in Foster et al. (2022), where it was concluded IUPAC ambiguity codes were beneficial for quality control, we conclude these may signal laboratory contamination (6). They may also indicate epidemiological importance where true iSNVs arise from host-to-host transmission or co-infection (27,28). Based on our observations, no universal minimum frequency threshold would have prevented the calling of a concordant iSNV which could be the result of participants receiving uneven allele mixtures or sequencing bias. Still, a good practice could be to call two consensus genomes using: 1) a strict minimum variant frequency threshold with ambiguity codes for the primary purpose of quality monitoring (i.e. a consensus sequence control); and 2) a major allele threshold without ambiguity codes for downstream analysis.

### Living guidelines for quality assessment during pandemics

Since the PTP analysed here concluded in 2021, several laboratory and bioinformatic developments have occurred, reflecting the evolving nature of the emergent pandemic response. The ARTIC primer scheme was redesigned in terms of laboratory methods and commercial schemes were offered to account for increasing variation (30,31). From a bioinformatic perspective, AusTrakka increased its minimum genome coverage criteria from >50% to >=90% (ACGT bases), which was the basis for the internal quality criterium of several participants. Regardless of AusTrakka participation, we agree with the conclusions of Lau et al. (2022) (13) and recommend this criterion as a minimum given the shared characteristics of the discordant metrics observed here (LB01-BS08, LB10-BS03, and LB10-BS08) being genome coverage <80% in addition to previous benchmarking studies (8,21). These developments support the need for living guidelines to continually review and update the quality standards for SARS-CoV-2 WGS.

### Value of the WG to Australian PHLs

A key advantage of the WG was the real-time collaboration and knowledge-sharing to form living guidelines by continually reviewing and updating the quality standards for SARS-CoV-2 WGS. One member noted that by comparing results across laboratories and methodologies, they could identify and rectify a spurious SNV impacting tree topology, which stemmed from a specific primer set. Another highlighted the utility of the Slack channel, finding it valuable for accessing bioinformatic tools shared by colleagues, thereby enriching their analytical processes. Amid the pressing challenges of pandemic surges on PHLs, this collaborative approach promoted bioinformatic equity via shared expertise and distributed tasks. Additionally, it offered a dynamic platform for discussing the continuously evolving understanding and demands associated with the virus.

### Limitations

Given this was a retrospective analysis, we must consider that the PTP was not designed for the study at hand. While it offered a dataset reflecting SARS-CoV-2 WGS methodologies across Australia, its design posed challenges for our comparative analysis. Issues included data entry mistakes and inconsistencies in the PTP questionnaire responses. The questionnaire aimed to gauge the ability of participants to accurately report and convey results, but our analysis highlights potential challenges PHLs faced during pandemic surges. This might also underscore varying PHL capabilities in recognizing critical bioinformatic traits. For future reference, multidisciplinary working groups might consider creating an intuitive bioinformatic analysis platform for participants to upload data and obtain technical feedback, like the metrics described here. These insights would increase bioinformatic analysis equity and provide PHLs with data-driven strategies to inform living guidelines, facilitating sequence data concordance as knowledge and methods evolve throughout a pandemic.

## Conclusions

Despite the variety of laboratory and bioinformatic methods used for SARS-CoV-2 WGS, the results were largely accurate. However, challenges such as low read depths and inconsistent amplicon sequencing coverage were observed, and variants at positions with read depths below 10 require careful consideration. We recommend PHLs bioinformatically generate two consensus genomes for distinct purposes: 1) quality control with ambiguity codes and a strict frequency threshold; and 2) downstream analysis without ambiguity codes. We also suggest implementing the following assessment criteria: 1) per-amplicon read depth as an indicator of primer efficiency; 2) SNVs to consider intrahost variation; and 3) a minimum genome coverage of 90%. In a genomics-forward era of public health pathogen surveillance, our study emphasizes the role of multidisciplinary working groups in providing technical analysis and feedback for sequence data concordance across PHLs. We also present additional assessment criteria to guide laboratories in identifying areas for improvement.

## Author contributions

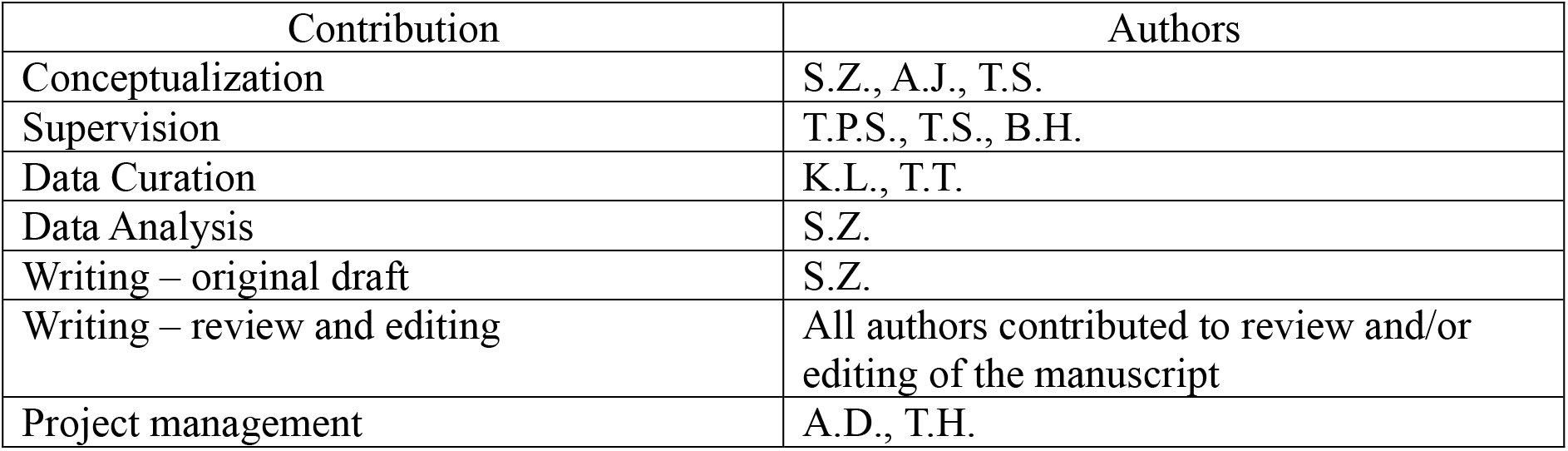

## Ethical approval

Human samples were not Primary samples from which the isolates were derived were collected in accordance with the Victorian Public Health and Wellbeing Act 2008. Ethical approval was received from the University of Melbourne Human Research Ethics Committee (study number 1954615.3).

## Acknowledgements

We acknowledge members of the MRFF COVID-19 Quality Control Working Group for their feedback and contribution to this paper. Members include: Jaisy Arikkatt, Susan Ballard, Angela Donald, Charles Foster, Rikki Graham, Tuyet Hoang, Amy Jennison, Jessica Johnson-Mackinnon, Jurissa Lang, Terrence Lee, Lex Leong, Jaimie McMahon, Paran Rayan Samuel, Torsten Seemann, Chisha Sikazwe, and Chenwei Wang. We also owe appreciation to the Royal College of Pathologists of Australasia Quality Assurance Programs (RCPAQAP), and The Communicable Diseases Genomics Network, supported by the Office of Health Protection and Response, within the Australian Government Department of Health and Aged Care, which has led the national coordination and integration of SARS-CoV-2 genomic data in Australia.

## References

1. Chen Z, Azman AS, Chen X, Zou J, Tian Y, Sun R, et al. Global landscape of SARS-CoV-2 genomic surveillance and data sharing. Nat Genet [Internet]. 2022 Apr;54(4):499x–507. Available from: 10.1038/s41588-022-01033-y

2. Lane CR, Sherry NL, Porter AF, Duchene S, Horan K, Andersson P, et al. Genomics-informed responses in the elimination of COVID-19 in Victoria, Australia: an observational, genomic epidemiological study. Lancet Public Health [Internet]. 2021 Aug;6(8):e547–56. Available from: 10.1016/S2468-2667(21)00133-X

3. Issues with SARS-CoV-2 sequencing data [Internet]. Virological. 2020 [cited 2022 Nov 15]. Available from: https://virological.org/t/issues-with-sars-cov-2-sequencing-data/473

4. Freed NE, Vlková M, Faisal MB, Silander OK. Rapid and inexpensive whole-genome sequencing of SARS-CoV-2 using 1200 bp tiled amplicons and Oxford Nanopore Rapid Barcoding. Biol Methods Protoc [Internet]. 2020 Jul 18;5(1):bpaa014. Available from: 10.1093/biomethods/bpaa014

5. Quick J, Grubaugh ND, Pullan ST, Claro IM, Smith AD, Gangavarapu K, et al. Multiplex PCR method for MinION and Illumina sequencing of Zika and other virus genomes directly from clinical samples. Nat Protoc [Internet]. 2017 Jun;12(6):1261–76. Available from: 10.1038/nprot.2017.066

6. Foster CSP, Stelzer-Braid S, Deveson IW, Bull RA, Yeang M, Au J-P, et al. Assessment of Inter-Laboratory Differences in SARS-CoV-2 Consensus Genome Assemblies between Public Health Laboratories in Australia. Viruses [Internet]. 2022 Jan 19;14(2). Available from: 10.3390/v14020185

7. Liu T, Chen Z, Chen W, Chen X, Hosseini M, Yang Z, et al. A benchmarking study of SARS-CoV-2 whole-genome sequencing protocols using COVID-19 patient samples. iScience [Internet]. 2021 Aug 20;24(8):102892. Available from: 10.1016/j.isci.2021.102892

8. Xiaoli L, Hagey JV, Park DJ, Gulvik CA, Young EL, Alikhan N-F, et al. Benchmark datasets for SARS-CoV-2 surveillance bioinformatics. PeerJ [Internet]. 2022 Sep 5;10:e13821. Available from: 10.7717/peerj.13821

9. Rambaut A, Holmes EC, O’Toole Á, Hill V, McCrone JT, Ruis C, et al. A dynamic nomenclature proposal for SARS-CoV-2 lineages to assist genomic epidemiology. Nat Microbiol [Internet]. 2020 Nov;5(11):1403–7. Available from: 10.1038/s41564-020-0770-5

10. Frampton D, Rampling T, Cross A, Bailey H, Heaney J, Byott M, et al. Genomic characteristics and clinical effect of the emergent SARS-CoV-2 B.1.1.7 lineage in London, UK: a whole-genome sequencing and hospital-based cohort study. Lancet Infect Dis [Internet]. 2021 Sep;21(9):1246–56. Available from: 10.1016/S1473-3099(21)00170-5

11. Geoghegan JL, Ren X, Storey M, Hadfield J, Jelley L, Jefferies S, et al. Genomic epidemiology reveals transmission patterns and dynamics of SARS-CoV-2 in Aotearoa New Zealand. Nat Commun [Internet]. 2020 Dec 11;11(1):6351. Available from: 10.1038/s41467-020-20235-8

12. Hoang T, da Silva AG, Jennison AV, Williamson DA, Howden BP, Seemann T. AusTrakka: Fast-tracking nationalized genomics surveillance in response to the COVID-19 pandemic. Nat Commun [Internet]. 2022 Feb 14;13(1):865. Available from: 10.1038/s41467-022-28529-9

13. Lau KA, Horan K, Gonçalves da Silva A, Kaufer A, Theis T, Ballard SA, et al. Proficiency testing for SARS-CoV-2 whole genome sequencing. Pathology [Internet]. 2022 Aug;54(5):615–22. Available from: 10.1016/j.pathol.2022.04.002

14. Lemoine F, Gascuel O. Gotree/Goalign: toolkit and Go API to facilitate the development of phylogenetic workflows. NAR Genom Bioinform [Internet]. 2021 Sep;3(3):qab075. Available from: 10.1093/nargab/lqab075

15. Quick J. nCoV-2019 sequencing protocol v3 (LoCost). protocols io. 08 2020;

16. Tyson JR, James P, Stoddart D, Sparks N, Wickenhagen A, Hall G, et al. Improvements to the ARTIC multiplex PCR method for SARS-CoV-2 genome sequencing using nanopore. bioRxiv [Internet]. 2020 Sep 4; Available from: 10.1101/2020.09.04.283077

17. Freed N, Silander O. SARS-CoV2 genome sequencing protocol (1200bp amplicon “midnight” primer set, using Nanopore Rapid kit). protocols io. 07 2021;

18. Eden J-S, Rockett R, Carter I, Rahman H, de Ligt J, Hadfield J, et al. An emergent clade of SARS-CoV-2 linked to returned travellers from Iran. Virus Evol [Internet]. 2020 Jan;6(1):veaa027. Available from: 10.1093/ve/veaa027

19. Eden J-S, Sim E. SARS-CoV-2 Genome Sequencing Using Long Pooled Amplicons on Illumina Platforms. protocols io. 04 2020;

20. O’Toole Á, Scher E, Underwood A, Jackson B, Hill V, McCrone JT, et al. Assignment of epidemiological lineages in an emerging pandemic using the pangolin tool. Virus Evol [Internet]. 2021 Jul 30;7(2):veab064. Available from: 10.1093/ve/veab064

21. Lau KA, Gonçalves da Silva A, Theis T, Gray J, Ballard SA, Rawlinson WD. Proficiency testing for bacterial whole genome sequencing in assuring the quality of microbiology diagnostics in clinical and public health laboratories. Pathology [Internet]. 2021 Dec;53(7):902–11. Available from: 10.1016/j.pathol.2021.03.012

22. Moran-Gilad J, Sintchenko V, Pedersen SK, Wolfgang WJ, Pettengill J, Strain E, et al. Proficiency testing for bacterial whole genome sequencing: an end-user survey of current capabilities, requirements and priorities. BMC Infect Dis [Internet]. 2015 Apr 3;15:174. Available from: 10.1186/s12879-015-0902-3

23. Gargis AS, Kalman L, Lubin IM. Assuring the Quality of Next-Generation Sequencing in Clinical Microbiology and Public Health Laboratories. J Clin Microbiol [Internet]. 2016 Dec;54(12):2857–65. Available from: 10.1128/JCM.00949-16

24. Borcard L, Gempeler S, Terrazos Miani MA, Baumann C, Grädel C, Dijkman R, et al. Investigating the Extent of Primer Dropout in SARS-CoV-2 Genome Sequences During the Early Circulation of Delta Variants. Frontiers in Virology [Internet]. 2022;2. Available from: https://www.frontiersin.org/articles/10.3389/fviro.2022.840952

25. Lambisia AW, Mohammed KS, Makori TO, Ndwiga L, Mburu MW, Morobe JM, et al. Optimization of the SARS-CoV-2 ARTIC Network V4 Primers and Whole Genome Sequencing Protocol. Front Med [Internet]. 2022 Feb 17;9:836728. Available from: 10.3389/fmed.2022.836728

26. Cotten M, Lule Bugembe D, Kaleebu P, T Phan M V. Alternate primers for whole-genome SARS-CoV-2 sequencing. Virus Evol [Internet]. 2021 Jan;7(1):veab006. Available from: 10.1093/ve/veab006

27. Armero A, Berthet N, Avarre J-C. Intra-Host Diversity of SARS-Cov-2 Should Not Be Neglected: Case of the State of Victoria, Australia. Viruses [Internet]. 2021 Jan 19;13(1). Available from: 10.3390/v13010133

28. Lythgoe KA, Hall M, Ferretti L, de Cesare M, MacIntyre-Cockett G, Trebes A, et al. SARS-CoV-2 within-host diversity and transmission. Science [Internet]. 2021 Apr 16;372(6539). Available from: 10.1126/science.abg0821

29. Davis JJ, Long SW, Christensen PA, Olsen RJ, Olson R, Shukla M, et al. Analysis of the ARTIC Version 3 and Version 4 SARS-CoV-2 Primers and Their Impact on the Detection of the G142D Amino Acid Substitution in the Spike Protein. Microbiol Spectr [Internet]. 2021 Dec 22;9(3):e0180321. Available from: 10.1128/Spectrum.01803-21

30. Bei Y, Pinet K, Vrtis KB, Borgaro JG, Sun L, Campbell M, et al. Overcoming variant mutation-related impacts on viral sequencing and detection methodologies. Front Med [Internet]. 2022 Oct 28;9:989913. Available from: 10.3389/fmed.2022.989913

