## Supplemental Figure 1 for "Bioinformatic investigation of discordant sequence data for SARS-CoV-2: insights for robust genomic analysis during pandemic surveillance"

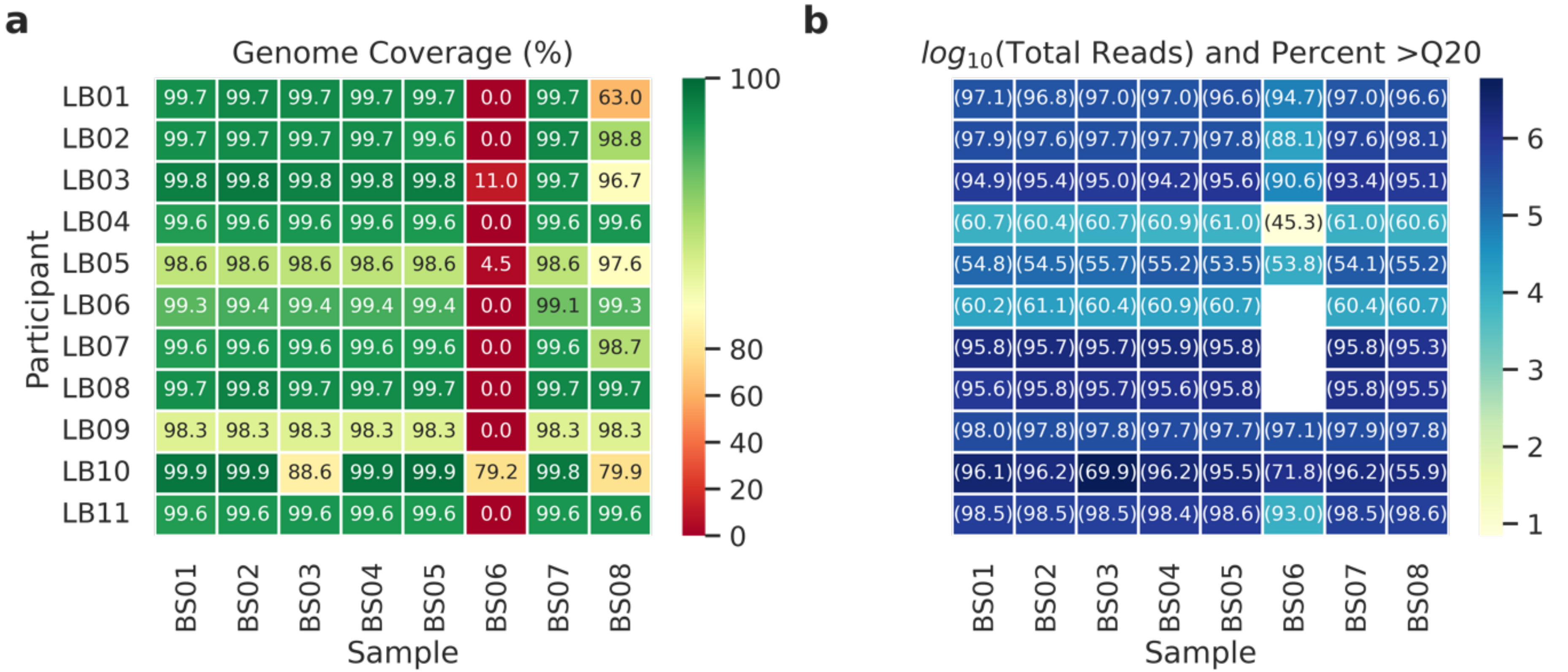


Supplementary Figure 1.

Characteristics of consensus genomes and reads submitted to the PTP. a) Genome coverage of consensus genomes submitted relative to the SARS-CoV-2 reference sequence (NC_045512.2). b) Number of reads per sample (log normalised), annotations indicate the percentage reads with quality score >Q20.
