## Supplemental Figure 2 for "Bioinformatic investigation of discordant sequence data for SARS-CoV-2: insights for robust genomic analysis during pandemic surveillance"

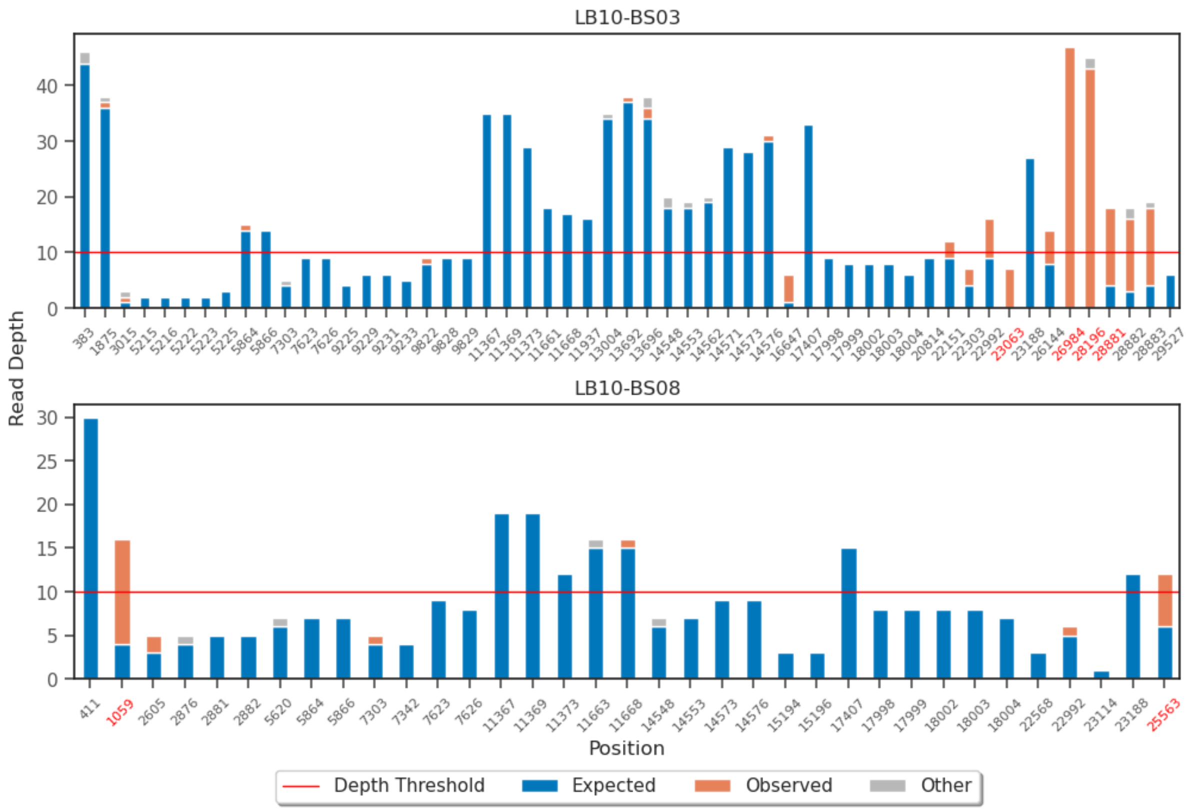


**Supplemental Figure 2.**

Allele frequencies at discordant positions from untrimmed reads of BS03 from LB10. The red horizontal line (Y=10) represents the generally accepted minimum read depth for variant calling for viral genomic surveillance. Nucleotide positions in red indicate the expected SNV sites.
